# Physical mechanisms that enable bacteria to traverse channels half their width

**DOI:** 10.1101/2025.03.07.642077

**Authors:** Kelsey G. DeFrates, Junsung Lee, Gissell Jimenez, Mariana G. Pinho, Christopher J. Hernandez

## Abstract

Bacteria colonize surfaces in the environment and can also penetrate tissues and materials by entering micro- and nanoscale cracks and pores. *Staphylococcus aureus* has been observed within nanoscale channels in bone that are 2-3 times smaller than cell diameter. Once inside bone, bacteria are protected from host immunity and systemic antibiotics, potentially contributing to chronic and recurrent infections. The physical mechanisms that enable bacteria to enter channels smaller than cell width are unclear. It has been proposed that bacteria traverse narrow passages through division – such that daughter cells form within small channels and eventually create a chain of cells extending down the channel length. Here we use microfluidics to test the idea that *S. aureus* can traverse bone-like nanochannels through growth. We examined 322 individual cells trapped within tapered nanochannels (∼1.5 to 0.4 µm in width). When cells were deformed below 606 nm (65% original width), growth and division were slowed or completely inhibited. When cell division did occur in nanochannels, daughter cells were more likely to travel towards the wider side of the channel. Hence, it is unlikely that cell division would preferentially enable transit of *S. aureus* into nanoscale channels. However, the magnitudes of fluid pressure needed to deform *S. aureus* to widths similar to that seen in bone (1 to 6.5 kPa), were small relative to fluid pressure in bone generated by physical forces *in vivo* (8-20 kPa). Thus, our findings suggest that colonization of nanoscale channels in bone or other tissues is more likely due to moderate fluid pressure rather than growth.

**Importance:** Bacteria that colonize materials and tissues within the body can be difficult to remove, even with thorough cleaning and application of antibiotics. Recent studies show that bacteria not only colonize the surfaces of tissues in the body but can also squeeze into naturally occurring pores and channels and thereby gain protection from immune cells and antibiotics. Here we ask how physical forces and cell growth might enable bacteria to enter small pores within materials. We use microfluidic devices to study the growth and migration of the human pathogenic bacteria, *S. aureus*, which is the leading cause of chronic bone infections.

## Main

Regulating bacterial colonization of materials and biological tissues is a significant challenge in human health. Bacterial infections of bone are among the most challenging to eradicate and require treatment by surgical debridement of the infection site and weeks of intravenous antibiotics (1). Despite the aggressive treatment regimen, 20% of cases show no improvement and rates of reinfection remain as high as 40% (2, 3). To improve therapeutic outcomes, a greater understanding of why current strategies fail is needed.

Recent studies suggest that recurrent infections in bone may be caused by the ability of pathogenic bacteria to colonize microarchitecture in the bone extracellular matrix (4–8). In cortical bone, fluid-filled channels run throughout the extracellular matrix to deliver nutrients to resident bone cells that are hundreds of micrometers away from vascular canals (**Fig. 1A**). These channels, called canaliculi, range in width from 300 to 900 nm (9). Several studies have observed *S. aureus* within the canalicular network of human and murine bone where cells are deformed to 1/3 their original width (**Fig. 1B**) (4–8). The ability of bacteria to colonize canaliculi is believed to contribute to infection persistence because neutrophils are too large to enter canaliculi and phagocytose resident bacteria. Additionally, the diffusion of systemic antibiotics into the canalicular space is limited, potentially contributing to bacterial survival (4–6).

**Figure 1.**
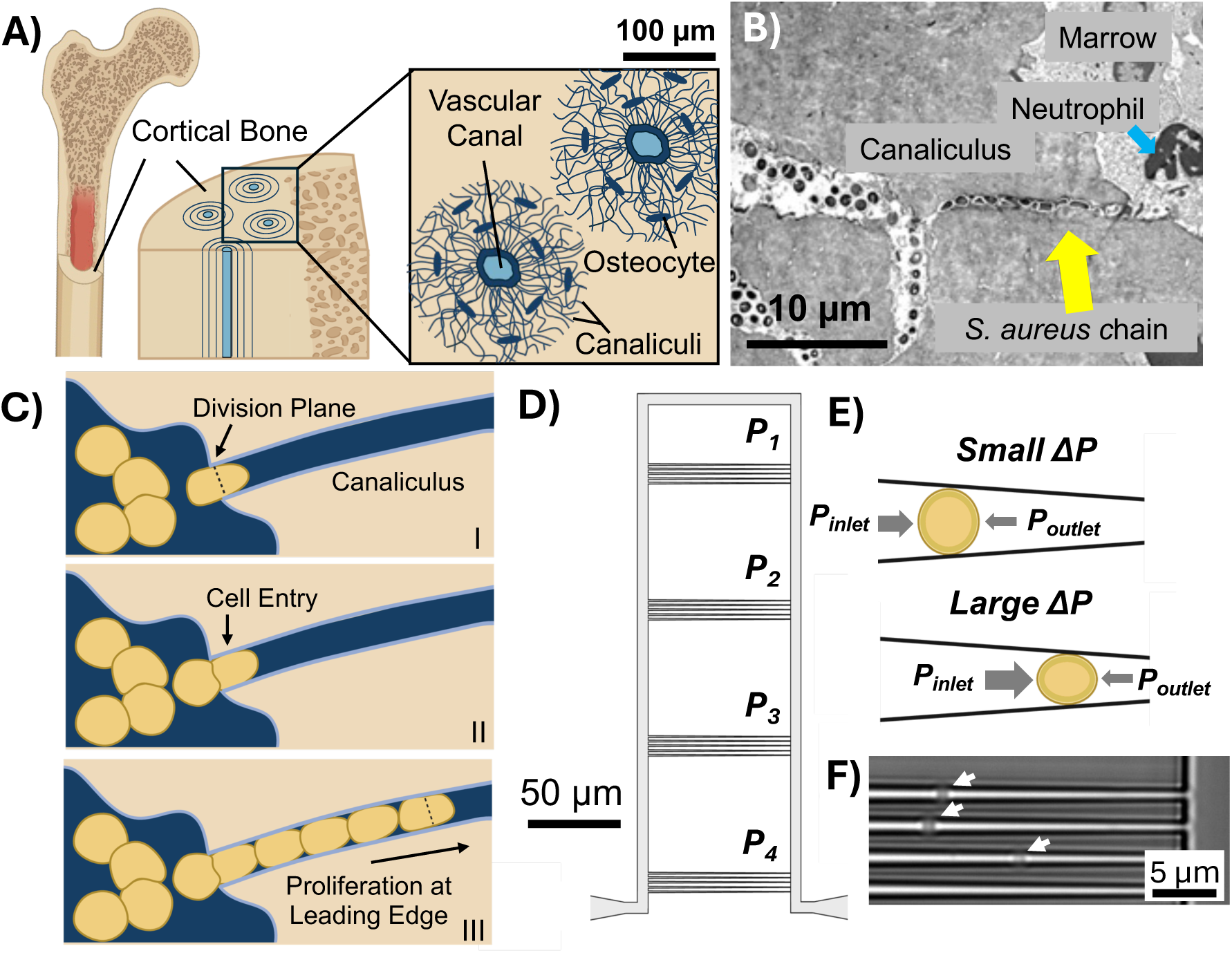
Schematic of the canalicular network in cortical bone is shown (A). Transmission electron microscopy of bone from a murine model of infection reveals *S. aureus* in canaliculi (yellow arrow) where cells are protected from neutrophils (blue arrow). From ref. 4 used with permission (B). Illustration of proposed mechanism of entry of *S. aureus* into bone through growth (C). The septal plane forms perpendicular to the channel opening (I), propelling one daughter cell into the canaliculus (II). Inside the channels, cells continue to divide leading to migration down the channel length (III). The extrusion loading device is shown with tapered channels connected in parallel to generate 4 distinct pressure levels (P_x_) for cell loading (D). The distance cells travel into the channels and the applied mechanical deformation is dependent upon the pressure differential (ΔP) generated between the channel inlet and outlet (E). Representative image of *S. aureus* cells (white arrows) within tapered channels is shown (F).

It is unclear how *S. aureus* traverses canaliculi, given that the cells are non-motile and 2 to 3 times larger than the width of the channels. Microfluidic systems have been useful for studying the transit of bacteria within nanoscale channels (10–13). Although motile bacteria can readily swim through channels that are only marginally larger than their diameter, bacteria may also enter channels smaller than their cell width through division. Migration through division is achieved when a single bacterium becomes lodged in a channel opening and divides to eventually give rise to a chain of growing daughter cells that extend the length of the channel (10). Through cell division, *Escherichia coli* (undeformed width 0.8 µm) was shown to navigate channels as small as 0.4µm in width. The ability of cells to migrate and grow within small spaces appears to be limited by the mechanical properties of the cell envelope, since the stiffer *Bacillus subtilis (*undeformed width 0.9 um) was unable to divide through passages smaller than 0.75 µm (10, 14). It has been proposed that *S. aureus* can also enter canaliculi in bone through cell division (4). To facilitate migration alignment of the septal plane across the channel width may anchor growing cells, leading to daughters cells being propelled forward into the nanochannel (**Fig. 1C**) (4). Evidence suggest that *S. aureus* may also preferentially sense and divide into canaliculi through a process known as durotaxis (4, 8, 15). In support of this hypothesis, *S. aureus* was found to propagate through 0.5 µm pores within a 0.4 µm thick silicon membrane (4, 8, 15). However, a thin, porous membrane does not recapitulate the channel-like geometry of canaliculi or allow for repeated cell divisions which would be needed to traverse canaliculi. Therefore, in this study we used microfluidic systems to better replicate the geometry of canaliculi and investigate *S. aureus* migration and growth within nanochannels.

We used a microfluidic device initially developed by our laboratory to study the biomechanics and mechanobiology of bacteria (14, 16–18). The device, which we call the ‘extrusion loading device,’ is manufactured from silica glass using deep UV lithography and consists of a series of tapered channels that narrow in width from ∼1.5 µm to 0.4 µm (**Fig 1D**). When a suspension of bacteria is flowed into the device, individual bacteria are trapped within the tapered channels by a pressure differential (ΔP) generated between the channel inlet and outlet (**Fig 1E**) (18). Greater pressure differences cause cells to travel further into the channel and experience greater deformation (**Fig. 1E**). To subject bacteria to a range of deformations, 4 distinct loading pressures are generated within one device by connecting sets of closely spaced channels in parallel (**Fig 1E**). Once inside the channels, prior work has verified that bacteria receive sufficient nutrients to support growth and viability for several hours (18). Here, we used the extrusion loading device to study deformation and growth of the methicillin-resistant strain of *S. aureus* (COL strain) within canaliculi-sized channels (**Fig 1F**). Across three biological replicate experiments, we analyzed 322 cells and applied differential pressure to achieve deformation of cells to canaliculi-like widths. To test the possibility of directional growth as a mechanism of *S. aureus* migration through bone, we determined the effects of mechanical deformation on the rate of cell division. We also determined if growth and division could result in net migration of daughter cells and whether cells showed a preference to divide downstream (toward the thinner side of the tapered channel) or upstream (towards the wider end of the channel).

Published work using the extrusion device found that *S. aureus* deformation inside submicron channels is proportional to the applied pressure allowing us to determine the mechanical properties (Young’s modulus) of the bacterial cell envelope through finite element modeling (19). In this study, we applied a limited range of differential fluid pressures from 1 to 6.5 kPa, which was anticipated to achieve cell deformation to canaliculi-like widths. To determine the original, undeformed cell width of *S. aureus*, we analyzed cells inside the extrusion loading device that were not transported into tapered channels and instead remained within feeder channels which measured several hundreds of microns in width. The average undeformed cell width was found to be 931 ± 41nm (**Fig S1**). In contrast, cells that were successfully loaded into tapered channels had widths ranging from 955 to 460 nm due to deformation by the channel walls (**Fig. 2A**). In alignment with our prior work with the device, we observed significant variability in deformed cell width at a given applied pressure (**Fig. S2**) (14, 16–18). We attribute this experimental variability to differences in starting cell width, the orientation of the Z ring relative to the channel length and/or the biomechanical properties of the cell. However, overall, we find that at all applied fluid pressures, *S. aureus* cells readily deformed within channels down to canaliculi-like widths (**Fig. S2**) (4).

**Figure 2.**
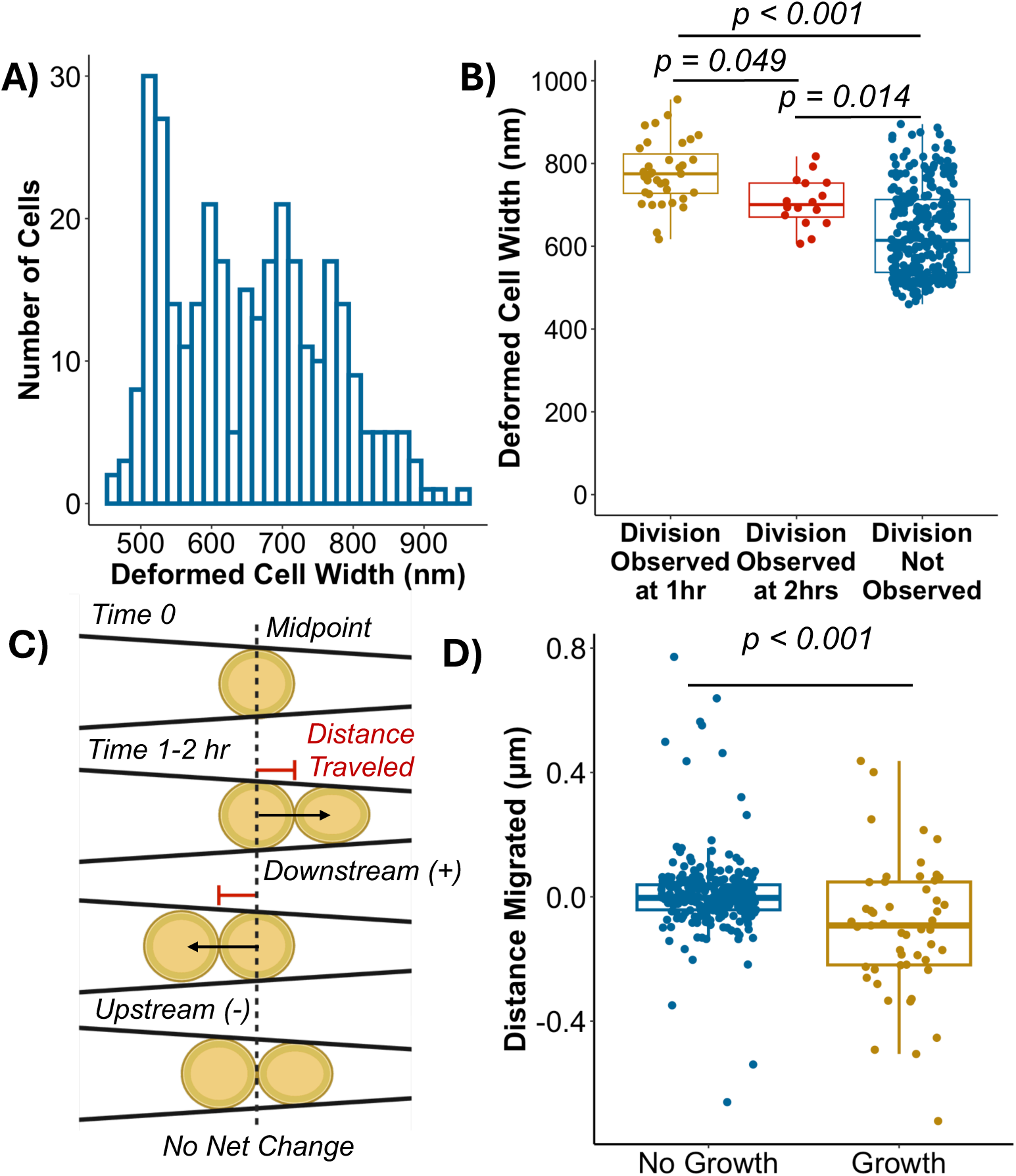
The distribution of deformed cell width for *S. aureus* after loading into the extrusion loading device is shown (N = 322 cells from 3 replicate experiments) (A). The correlation between deformed cell width and time to division is shown (B). Cells that were found to divide during the first interval hour after loading within the microfluidic device had a larger cell width (778 *±* 77nm, mean ± sd n = 35) than those that divided during the second hour (706 *±* 59nm, n = 16), or did not divide at all over the 3 hr observation period (634 *±* 104nm, n = 271). Schematic illustration of analysis method used to determine the distance migrated after division (C). The distance migrated for cells that divided in hour 1 or 2 of the experiment (n = 51) and for cells that did not divide (n = 271) is shown. Statistics are based on a One-way ANOVA with Tukey HSD for (A) and Student’s t-test for (D).

To determine if directional growth plays a role in *S. aureus* transport into canaliculi, we investigated the effects of mechanical confinement inside the extrusion loading device on cell division. We observed an effect of cell deformation on time to division: cells that divided during the first hour of the experiment tended to be less deformed than cells that divided by the second hour (deformed cell width 778 ± 77nm vs. 706 ± 59nm) (**Fig 2B**). Cells that did not divide at all over the course of the experiment were also generally subjected to the most significant degree of deformation (634 ± 104nm) and no divisions were detected in cells measuring less than 606nm in width (maximum observation period 3 hrs, **Fig. 2B**). Thus, mechanical confinement appears to slow cell division and deformation below ∼65% cell width severely inhibits *S. aureus* growth.

If *S. aureus* traverses canaliculi through growth and preferentially divides into bone, we hypothesized that cell division inside the extrusion loading device would result in net migration of cells towards the narrower end of the channel. Therefore, when division occurred, we determined whether daughter cells migrated by measuring the change in midpoint between the original cell and daughter cell pair (**Fig 2C**). We found that growth and division had a marginal effect on cell position within the channel. On average, cells migrated only 97 ± 215nm and showed a slight bias (56% of cells studied) to migrate upstream towards the wider end of the channel (**Fig 2D**).

Together with our observation of impaired growth in deformed cells, we conclude that *S. aureus* is unlikely to divide into and grow through canaliculi that measure less than ∼600 nm in width. Our results suggest that it is more likely that *S. aureus* enters canaliculi when cells are forced into bone by fluid pressure. We found that differential fluid pressures as low as 1 to 6.5 kPa are sufficient to force *S. aureus* into spaces that are nearly half pf cell diameter. This pressure magnitude is substantially less than the fluid pressures measured at the surface of bone during normal movement (8 to 20 kPa) (20). Although *S. aureus* are stiff, Gram-positive organisms that may not be expected to deform substantially under fluid pressure, recent findings show that the stiffness (Young’s modulus) of the cell envelope of *S. aureus* (1.52 ± 0.06 MPa) is of similar magnitude to that of Gram-negative *E. coli* (2.06 ± 0.04 MPa) (16). Thus, the ability of bacteria to colonize bone may be more closely related to physical loads on the bone during activity and the mechanical properties of the bone matrix and bacteria rather than growth processes.

In addition to elucidating the mechanism by which *S. aureus* invades nanocavities in bone, our work provides insight into how bacteria may be transported through different microenvironments. We have shown that relatively low fluid pressures can force bacteria into spaces that are substantially smaller than cell width. Micro- and nanoscale cavities are ubiquitous in biological tissues and materials in the natural and built environment. Bacteria that colonize small spaces may be protected from predation, sterilization, and antimicrobial therapies, and may continue to divide if provided sufficient nutrients (21, 22). Purposefully encapsulating bacteria within materials to create a new class of composites known as ‘engineered living materials’ has also proven useful in creating structures with stimuli-responsive behaviors, self-healing properties, and therapeutic utility (23–25). Our study demonstrates how it may be possible to transport bacteria into synthetic scaffolds or isolated areas of the human body using fluid pressure.

## Methods

### Cell culturing

*S. aureus* COL strain (NRS100) was grown overnight (18 hrs) in Tryptic Soy Broth (TSB) (Millipore Sigma) at 37 °C with 200 rpm shaking. The overnight culture was diluted 1:200 in TSB and grown for 3 hrs at 37 °C until OD600 ∼ 0.3 to reach exponential phase.

### Microfluidic device manufacturing

Microfluidic device manufacturing was done using Deep UV photolithography as previously described (14, 16–18). Briefly, fused silica wafers (100 mm diameter and 500 µm thick, WF3937X02031190, Mark Optics, Santa Ana, CA, USA) were coated with ∼ 55 nm of chrome using the AJA Sputter Deposition Tool (AJA International, Scituate MA, USA). A ∼60 nm coat of anti-reflective coating (ARC, DUV 42P, Brewer Science, Rolla, MO, USA) and ∼ 510 nm coat of photoresist (UV210, MicroChem, Westborough, MA, USA) were then applied using The Gamma Automatic Coat-Develop Tool (Suss MicroTec Gamma Cluster Tool, Garching Germany). The custom microfluidic device pattern was transferred to the wafer using the ASML Deep UV stepper (Veldhoven Netherlands). The photoresist was then developed using the Gamma Automatic Coat-Develop Tool and the pattern was transferred from the photoresist to the anti-reflective coating using plasma etching in the Oxford 82 Tool (Oxford, Abingdon, UK). The pattern was transferred to the chrome layer using the Plasma-Therm 770 ICP tool (Plasma-Therm St. Petersburg FL, USA). Oxygen plasma cleaning by the Oxford 82 Tool was performed to remove residual coating before the pattern was finally transferred to the silica wafer using the Oxford 100 Tool (Oxford, Abingdon, UK). Any remaining chrome was removed using a wet chemical bath. Through-holes were laser-etched at the microfluidic device inlets and outlets using a Versalaser (VLS3.50, Universal Laser Systems, Scottsdale, AZ, USA) to establish inlet and outlet holes. All fabrication steps were performed within the cleanroom of the Cornell NanoScale Facility Science and Technology Facility (Ithaca, NY, USA).

To verify that the pattern was successfully transferred, device feature dimensions were characterized using atomic force microscopy (Veeco Icon Bruker, Billerica MA, USA), profilometry (P-7, KLA Inc, Milpitas CA, USA) and scanning electron microscopy (Zeiss Ultra 55 SEM microscope, Oberkocken Germany). Devices used in this study had an average depth of 1.01 ± 0.01 um, tapper inlet width of 1.73 ± 0.46 µm and outlet width of 0.46 ± 0.025 um.

Successfully patterned wafers were bonded to silica cover wafers (100 mm diameter and 170 µm thick) (WF3937X0073119B Mark Optics, Santa Ana CA, USA) after MOS/RCA cleaning by hand bonding and nitrogen annealing (5 hrs, 1100 °C).

### *S. aureus* within the microfluidic device

To load cells into the microfluidic device, fluid pressure was supplied by a PneuWave Pump (CorSolutions, Ithaca NY, USA). The pump was attached to the microfluidic device inlet using PEEK tubing (Idex 360 µm OD × 150 µm ID, Lake Forest IL, USA) that was feed through a magnetic connector lever arm (Fluidic Indexing Probe, CorSolutions, Ithaca NY, USA). A rubber gasket (N-123-03 IDEX, Lake Forest IL, USA) was also included at the end of the tubing to prevent leakage. Before loading cells, the tubing was sterilized by running 10% bleach followed by 70% ethanol through the pump for 15 min. TSB was then flowed through the tubing to waste for 20 minutes before attaching to the device. The device was wet with TSB for 30 minutes. The tubing was then disconnected from the device and the cell suspension was flushed through for 15 minutes into waste. The tubing was then reattached to the device and 60 or 80 kPa of pressure was applied to load cells within tapered channels. Two different loading pressures were used to ensure that the observed results were the effect of cell deformation by the channel walls, rather than flow rate or hydrostatic loading. Media was flowed continuously through the device for the remainder of the experiment. To observe cell loading and division, the device was mounted on an Olympus IX83 inverted microscope (Evident Scientific, Waltham MA, USA) equipped with a motorized stage (Märzhäuser Wetzlar SCAN IM, Wetzlar, Germany) and Okolab Cage Incubator set to 37°C (H201-Enclosure with temperature control unit, Okolab, Sewickley PA, USA). Immediately after loading and hourly for up to 3 hours, brightfield images were taken of cells in individual tappers using a 100x objective (UPLXAPO100X0, Evident Scientific, Waltham MA, USA).

### Measuring cell boundaries and migration within the microfluidic device

A custom Matlab (v. 2023b, Mathworks, Natick, MA, USA) script was developed to measure cell width and distance traveled within the microfluidic device (14, 16–18). The script requires user input to identify regions of interest within brightfield images containing cells. Horizontal and vertical line profiles are then taken within the region. The distance between the midpoints in curves from the horizontal and vertical profiles are used to estimate cell length and width, respectively (**Fig. S3**). Users then identify the channel end and the distance between the midpoint of the cell and outlet is calculated. The distance cells traveled in the channel was also used to calculate the expected deformed width based on the channel geometry. To determine the distance migrated overtime, the change in distance of cells in the taper between time 0 and time 1 (for cells that did not divide over the course of the entire experiment or cells that divided within the first hour), or time 1 and time 2 (for cells that divided in the second hour) was determined. When cells divided, the user defined regions of interest that included the daughter cell pair. The midpoint of the two cells combined was then compared to the midpoint of the original cell.

If cells did not divide over the course of the experiment, the change in distance for the single cell between time 1 and time 2 was determined. A frame of reference was established so that when cells moved downstream (towards the narrower end of the taper), this resulted in a net positive change in distance, while movement upstream (towards the wider end) resulted in a negative change. Tapers initially loaded with more than one cell were excluded from analysis.

To determine undeformed cells width, cells residing near the inlet of the microfluidic device that were not transported into tapered channels were imaged and average width was calculated using the MicrobeJ plugin on ImageJ.

### Microfluidic device hydraulic circuit pressure calculations

To determine the fluid pressure at the tapered channels within the microfluidic devices, we performed hydraulic circuit calculations using a custom script in MATLAB (v. 2023b, Mathworks, Natick, MA, USA) as described (14, 17). In the analysis, the Hagen-Poiseuille law (Eqn 1) is used to determine the pressure drop, *ΔP*, across each channel with flow rate *Q* and hydraulic resistance, *R_h_*.

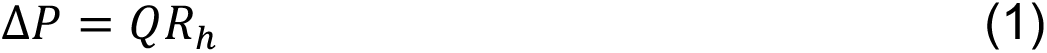

To calculate *R_h_*, Poiseuille flow (Eqn 2 and 3) was used when the ratio of the channel width to height was small (<20), while Plane Poiseuille flow (Eqn 4) was used when width/height was larger (>20). In each case, hydraulic resistance is a function of the fluid viscosity, *µ* (assumed water = 8.9e−4 Pa s), the length of the channel, *L*, the cross-sectional area of the channel, *A*, and the hydraulic radius, *r*. For a rectangular channel, *r* is calculated from the perimeter of the channel cross-section, *P*, and channel height, *H*.

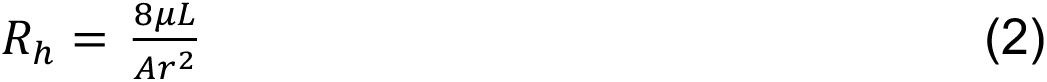

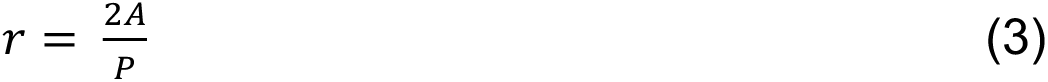

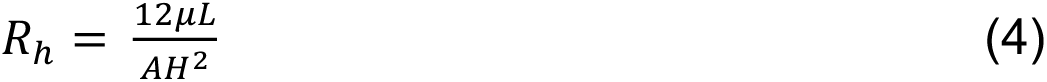

To determine *R_Total_*, the total hydraulic resistance across the device *R_h_* values for all channels were combined. When channels were connected in parallel, R_Total_ was calculated using Eqn 5 for *n* number of channels.

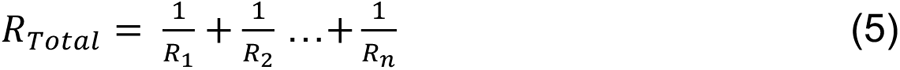

Channels connected in series were combined using Eqn 6.

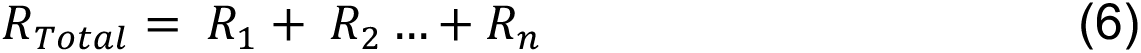

### Statistical Analysis

One-way ANOVA with post-hoc Tukey HSD or Student’s t-test was used to compare groups when appropriate. Statistical analyses were performed with a significance level of α = (0.05). Data were analyzed using RStudio.

## Acknowledgements

Financial support was provided by the NSF BRITE award (CMMI-2135586) and seed grant provided by the UCSF Center for Disruptive Musculoskeletal Innovation. This work was performed in part at the Cornell NanoScale Facility, a member of the National Nanotechnology Coordinated Infrastructure (NNCI), which is supported by the National Science Foundation (Grant NNCI-2025233). Biorender was used to make some figures.

## Author Contributions

K.G.D. wrote the manuscript, performed cell loading and growth experiments and imaging, analyzed and curated data, and performed statistical analysis. J.L assisted with experimental troubleshooting, manufactured devices, and image analysis. J.G. assisted with image analysis. M.G.P. contributed the cell strains, consulted on research design and data interpretation, and edited the manuscript. C.J.H. designed and directed research, edited manuscript.

## Data Availability

The data that support the findings of this study are available within the article and its supplementary material.

## References

1. Conterno LO, Turchi MD. 2013. Antibiotics for treating chronic osteomyelitis in adults. Cochrane Database Syst Rev CD004439.

2. Masters EA, Trombetta RP, de Mesy Bentley KL, Boyce BF, Gill AL, Gill SR, Nishitani K, Ishikawa M, Morita Y, Ito H, Bello-Irizarry SN, Ninomiya M, Brodell JD, Lee CC, Hao SP, Oh I, Xie C, Awad HA, Daiss JL, Owen JR, Kates SL, Schwarz EM, Muthukrishnan G. 2019. Evolving concepts in bone infection: redefining “biofilm”, “acute vs. chronic osteomyelitis”, “the immune proteome” and “local antibiotic therapy.” Bone Res 7:1–18.

3. Schwarz EM, Parvizi J, Gehrke T, Aiyer A, Battenberg A, Brown SA, Callaghan JJ, Citak M, Egol K, Garrigues GE, Ghert M, Goswami K, Green A, Hammound S, Kates SL, McLaren AC, Mont MA, Namdari S, Obremskey WT, O’Toole R, Raikin S, Restrepo C, Ricciardi B, Saeed K, Sanchez-Sotelo J, Shohat N, Tan T, Thirukumaran CP, Winters B. 2019. 2018 International Consensus Meeting on Musculoskeletal Infection: Research Priorities from the General Assembly Questions. Journal of Orthopaedic Research 37:997–1006.

4. de Mesy Bentley KL, Trombetta R, Nishitani K, Bello-Irizarry SN, Ninomiya M, Zhang L, Chung HL, McGrath JL, Daiss JL, Awad HA, Kates SL, Schwarz EM. 2017. Evidence of Staphylococcus Aureus Deformation, Proliferation, and Migration in Canaliculi of Live Cortical Bone in Murine Models of Osteomyelitis. Journal of Bone and Mineral Research 32:985–990.

5. 5. de Mesy Bentley KL, MacDonald A, Schwarz EM, Oh I. 2018. Chronic Osteomyelitis with Staphylococcus aureus Deformation in Submicron Canaliculi of Osteocytes: A Case Report. JBJS Case Connector 8:e8.

6. Zoller SD, Hegde V, Burke ZDC, Park HY, Ishmael CR, Blumstein GW, Sheppard W, Hamad C, Loftin AH, Johansen DO, Smith RA, Sprague MM, Hori KR, Clarkson SJ, Borthwell R, Simon SI, Miller JF, Nelson SD, Bernthal NM. 2020. Evading the host response: Staphylococcus “hiding” in cortical bone canalicular system causes increased bacterial burden. Bone Res 8:1–11.

7. Masters EA, Bentley KL de M, Gill AL, Hao SP, Galloway CA, Salminen AT, Guy DR, McGrath JL, Awad HA, Gill SR, Schwarz EM. 2020. Identification of Penicillin Binding Protein 4 (PBP4) as a critical factor for Staphylococcus aureus bone invasion during osteomyelitis in mice. PLOS Pathogens 16:e1008988.

8. Masters EA, Muthukrishnan G, Ho L, Gill AL, de Mesy Bentley KL, Galloway CA, McGrath JL, Awad HA, Gill SR, Schwarz EM. 2021. Staphylococcus aureus Cell Wall Biosynthesis Modulates Bone Invasion and Osteomyelitis Pathogenesis. Front Microbiol 12.

9. Marotti G, Ferretti M, Remaggi F, Palumbo C. 1995. Quantitative evaluation on osteocyte canalicular density in human secondary osteons. Bone 16:125–128.

10. Männik J, Driessen R, Galajda P, Keymer JE, Dekker C. 2009. Bacterial growth and motility in sub-micron constrictions. Proceedings of the National Academy of Sciences 106:14861–14866.

11. Takeuchi S, DiLuzio WR, Weibel DB, Whitesides GM. 2005. Controlling the Shape of Filamentous Cells of Escherichia coli. Nano Lett 5:1819–1823.

12. Figueroa-Morales N, Rivera A, Soto R, Lindner A, Altshuler E, Clément É. 2020. E. coli “super-contaminates” narrow ducts fostered by broad run-time distribution. Science Advances 6:eaay0155.

13. Lynch JB, James N, McFall-Ngai M, Ruby EG, Shin S, Takagi D. 2022. Transitioning to confined spaces impacts bacterial swimming and escape response. Biophysical Journal 121:2653–2662.

14. Sun X, Weinlandt WD, Patel H, Wu M, Hernandez CJ. 2014. A microfluidic platform for profiling biomechanical properties of bacteria. Lab Chip 14:2491–2498.

15. Masters EA, Salminen AT, Begolo S, Luke EN, Barrett SC, Overby CT, Gill AL, de Mesy Bentley KL, Awad HA, Gill SR, Schwarz EM, McGrath JL. 2019. An in vitro platform for elucidating the molecular genetics of S. aureus invasion of the osteocyte lacuno-canalicular network during chronic osteomyelitis. Nanomedicine 21:102039.

16. Lee J, Jha K, Harper CE, Zhang W, Ramsukh M, Bouklas N, Dörr T, Chen P, Hernandez CJ. 2024. Determining the Young’s Modulus of the Bacterial Cell Envelope. ACS Biomater Sci Eng 10:2956–2966.

17. Harper CE, Zhang W, Lee J, Shin J-H, Keller MR, van Wijngaarden E, Chou E, Wang Z, Dörr T, Chen P, Hernandez CJ. 2023. Mechanical stimuli activate gene expression via a cell envelope stress sensing pathway. Sci Rep 13:13979.

18. Genova LA, Roberts MF, Wong Y-C, Harper CE, Santiago AG, Fu B, Srivastava A, Jung W, Wang LM, Krzemiński Ł, Mao X, Sun X, Hui C-Y, Chen P, Hernandez CJ. 2019. Mechanical stress compromises multicomponent efflux complexes in bacteria. Proceedings of the National Academy of Sciences 116:25462–25467.

19. Lee J, Jha K, Harper CE, Zhang W, Ramsukh M, Bouklas N, Dörr T, Chen P, Hernandez CJ. 2024. Determining the Young’s Modulus of the Bacterial Cell Envelope. ACS Biomater Sci Eng 10:2956–2966.

20. Robertsson O, Wingstrand H, Kesteris U, Jonsson K, Onnerfält R. 1997. Intracapsular pressure and loosening of hip prostheses. Preoperative measurements in 18 hips. Acta Orthop Scand 68:231–234.

21. Wang P, Robert L, Pelletier J, Dang WL, Taddei F, Wright A, Jun S. 2010. Robust Growth of *Escherichia coli*. Current Biology 20:1099–1103.

22. Kalairaj MS, George I, George SM, Farfán SE, Lee YJ, Rivera-Tarazona LK, Wang S, Abdelrahman MK, Tasmim S, Dana A, Zimmern PE, Subashchandrabose S, Ware TH. 2024. Controlled release of microorganisms from engineered living materials. bioRxiv 10.1101/2024.09.25.615042.

23. Heveran CM, Hernandez CJ. 2023. Make engineered living materials carry their weight. Matter 6:3705–3718.

24. Rodrigo-Navarro A, Sankaran S, Dalby MJ, del Campo A, Salmeron-Sanchez M. 2021. Engineered living biomaterials. Nat Rev Mater 6:1175–1190.

25. Srubar WV. 2021. Engineered Living Materials: Taxonomies and Emerging Trends. Trends in Biotechnology 39:574–583.

